# Parallel shifts of visual sensitivity and body colouration in replicate populations of extremophile fish

**DOI:** 10.1101/2021.06.16.448734

**Authors:** Gregory L. Owens, Thor Veen, Dylan R. Moxley, Lenin Arias-Rodriguez, Michael Tobler, Diana J. Rennison

## Abstract

Visual sensitivity and body pigmentation are often shaped by both natural selection from the environment and sexual selection from mate choice. One way of quantifying the impact of the environment is by measuring how traits have changed after colonization of a novel habitat. To do this, we studied *Poecilia mexicana* populations that have repeatedly adapted to extreme sulphidic (H_2_S containing) environments. We measured visual sensitivity using opsin gene expression, as well as body pigmentation and water transmission for populations in four independent drainages. Both visual sensitivity and body pigmentation showed significant parallel shifts towards greater medium wavelength sensitivity and reflectance in sulphidic populations. The light spectrum was only subtly different between environments and overall, we found no significant correlations between the light environment and visual sensitivity or body pigmentation. Altogether we found that sulphidic habitats select for differences in visual sensitivity and pigmentation; our data suggest that this effect is unlikely to be driven purely by the water’s spectral properties and may instead be from other correlated ecological changes.

## Introduction

Patterns of parallel and convergent evolution are strong evidence of the action of natural selection, as it is unlikely that drift would lead to the same phenotype evolving in multiple independently derived populations or species (Schluter and Nagel 1995). Due to vision’s central role in predation avoidance, mate choice, and foraging, it is predicted to be under strong natural and/or sexual selection in many species (Endler 1992). Indeed, work in a variety of systems has indicated that shifts in visual system do evolve repeatedly (O’Quin et al. 2010; Rennison et al. 2016; Torres-Dowdall et al. 2017). These shifts have often been found to be largely genetically determined (e.g., (Tobler et al. 2010; Rennison et al. 2016)), although phenotypic plasticity also plays a role (e.g., (Nandamuri et al. 2017)). Yet, identification of the ecological factors and functional mechanisms shaping these patterns of evolution has proven difficult. The visual system is predicted to evolve to roughly match the availability of wavelengths to maximize photon catch and contrast detection through natural selection (Clarke 1957; Denton and Warren 1957; Munz 1958). But even in cases where there is some evidence of matching over portions of the visual spectrum, overall shifts in visual sensitivity remain largely unexplained by hypotheses related to background matching (e.g., (Rennison et al. 2016)).

Apart from natural selection, sexual selection may also be playing a role in determining visual sensitivity. Shifts in visual sensitivity are often accompanied by differences in body pigmentation and colour-based mate choice. The sensory bias and sensory drive hypotheses attempt to explain patterns of coevolution between male signals and female perception. These hypotheses suggest that male sexual signals should become tuned to match the sensitivity of the female’s sensory system to optimize attractiveness (Boughman 2002; Fuller et al. 2005). Sensory bias posits that mate signals should match sensory perception and is exemplified in taxa such as swordtail fish (Basolo 1990) and tungara frogs (Ryan and Rand 1990). In contrast, sensory drive integrates natural selection and proposes that while signals and perception should match, both are constrained and influenced by the environment. African cichlids (Seehausen et al. 2008) and threespine stickleback (Boughman 2001) are among the few systems where sensory drive seems to explain patterns of co-evolution between shifts in female visual perception and male nuptial colouration. In general, few studies have described a pattern of coevolution between visual perception and body pigmentation (but see (Brock et al. 2018)) or determined the pattern of evolution of each trait in relation to ambient light. Thus, it remains unclear what ecological mechanisms most often drive shifts in body pigmentation and visual sensitivity. Further, when signalling conditions seem relatively benign, for example, in a habitat where ambient light is generally broad spectrum and/or differences in ambient light are subtle between habitats, it’s unclear whether the selective pressure is strong enough to drive spectral matching.

*Poecilia* fish inhabiting sulphide springs in Mexico are a phenomenal example of convergent evolution (Tobler et al. 2018). These fish have evolved to survive in the presence of hydrogen sulphide (H2S), a potent respiratory toxicant (Tobler et al. 2016) and adaptation has been repeated in multiple independently colonized locations (Tobler et al. 2011). Sulphidic and non-sulphidic populations have been documented to diverge in physiological (Greenway et al. 2020), morphological (Tobler and Hastings 2011) and life history traits (Riesch et al. 2010b). In addition, populations in adjacent sulphidic and non-sulphidic habitats are reproductively isolated and exhibit very low levels of gene flow despite a lack of physical barriers that would prevent fish movement (Plath et al. 2013). Aside from the presence of H2S, the colonized habitats also vary in other ecological properties compared to the ancestral non-sulphidic habitats, including the availability of food resources (Tobler et al. 2015) and community composition (presence of predators & competitors) 16/06/2021 15:40:00.

Sulphur containing solutions (aqueous and non-aqueous) are known to absorb wavelengths in the ultraviolet (200-360 nm) region (Okada 1963; Khan 2011). This suggests the ambient light environment may also differ between the adjacent sulphidic and non-sulphidic locations and may drive shifts in visual sensitivity and/or body pigmentation. We surveyed four independently colonized drainages with paired sulphidic and non-sulphidic sites containing *Poecilia* species to ask the following questions: 1) Has there been parallel evolution of shifts in visual sensitivity and/or body pigmentation of the sulphidic and non-sulphidic ecotypes across the different drainages? 2) Has there been co-evolution between female perception and body pigmentation? 3) Are there differences in the light environment between the habitats that explain any observed phenotypic shifts?

## Methods

### Sample collection

Specimens of *Poecilia mexicana* were collected from four drainages in the Río Grijalva basin (from west to east: Pichucalco, Ixtapangajoya, Puyacatengo, and Tacotalpa). In each drainage, we sampled fish from one sulphidic (La Gloria springs, La Esperanza springs, La Lluvia springs, El Azufre) and one non-sulphidic (Rio El Azufre west branch, Rio Ixtapangajoya, Rio Puyacatengo at Vicente Guerrero, and Arroyo Bonita) population (Figure 1). Ten female individuals were sampled from each population and euthanized using MS222 for opsin expression analysis. Reflectance measurements were taken from 10-15 live male and female fish from each location (30 fish total per population). During transport, the live fish were held in black buckets for one to two hours before spectral measurements. Three replicated measurements were taken from each of four body locations (top of head, behind the eye, abdomen, and tail). Due to technical constraints, we only measured reflectance in fully opaque body regions. Measurements taken at partially transparent body parts, for example fins, produced inconsistent measurements between replicates. This means that our tail measurement is of the caudal peduncle and not the caudal fin. At each of the eight locations we collected fish, we also measured the *in situ* spectral conditions from 351 to 700 nm. Irradiance measurements of side-welling light were taken at depths of 0, 10, 20 and 30 cm (maximal depth) at five or six sites within each sampling location using a cosine corrector attached to a spectrophotometer (Ocean Optics, USA).

**Figure 1.**
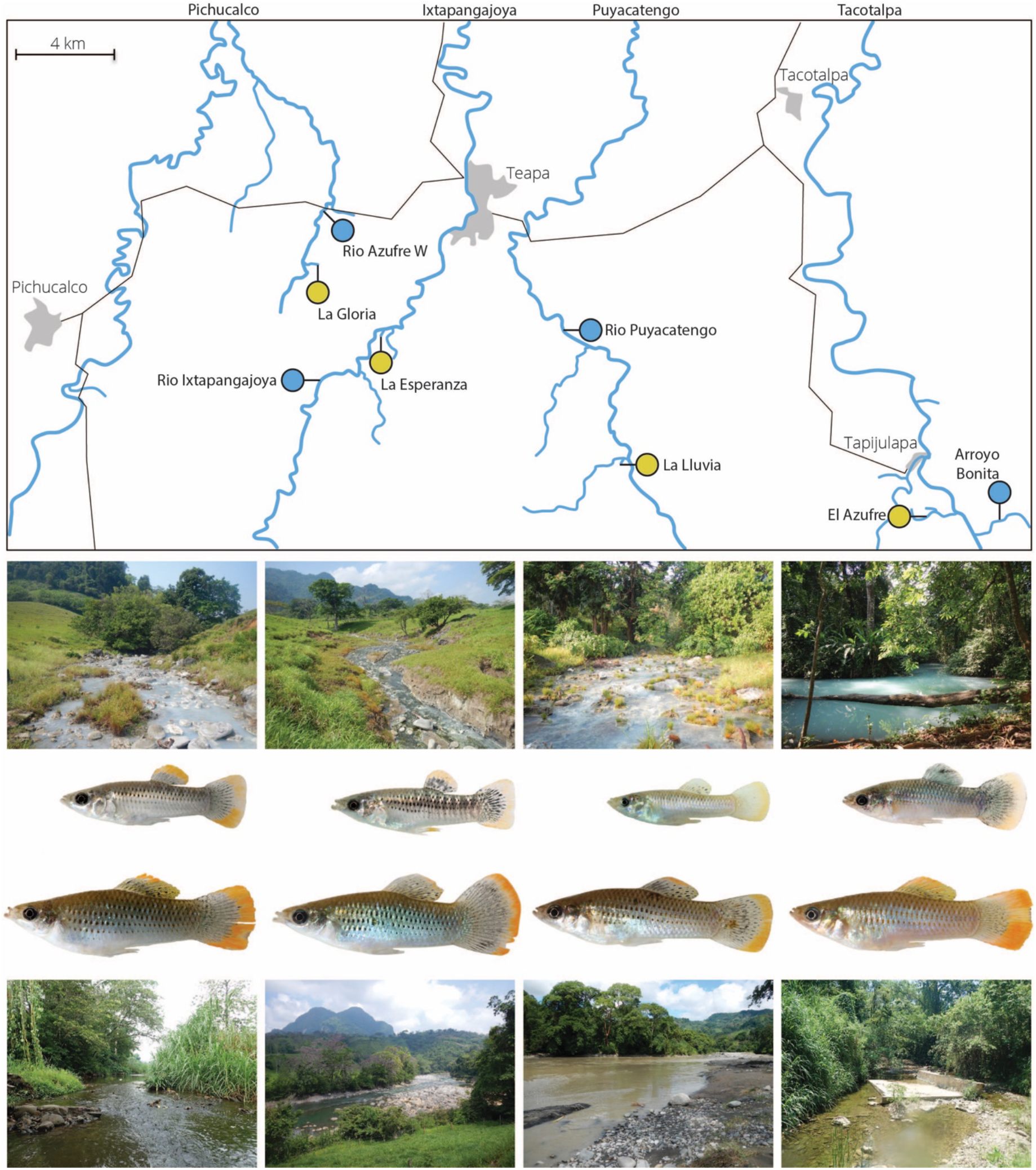
Map of the study region including the four drainages with paired sulphidic (yellow) and non-sulphidic (blue) sites. Photos show sulphidic habitats and *P. mexicana* (top row) non-sulphidic habitats and *P. mexicana* (bottom row) for each of the drainages.

### Estimation of environmental water light transmission

Following Rennison et al. (2016), we converted irradiance measurements to Einsteins, smoothed each measurement over a five nm rolling mean, and then fit a spline using the splinefun function in R (v3.5.1) (R Development Core Team 2013). Irradiance is a measure of current light conditions and varies with both water depth as well as the weather and angle of the sun. Therefore, we focused on transmission, which measures the amount of photon loss through water, is a property of the water, and should be more stable over short time scales. Our measure of transmission, K_S_, was calculated as the 1−A_S_, the light extinction coefficient with depth (see Rennison et al. 2016 for details). A transmission value of one indicates that no photons are being absorbed by the water, so the irradiance is identical at the surface and at lower depths. Lower values of transmission indicate less photons are transmitting through the water at that particular wavelength. For one sampling site in one location, we found transmission values > 1 across large regions of the spectrum suggesting technical problems, so we removed this data point.

### Opsin expression and spectral sensitivity

Both eyes were removed immediately after euthanasia, stored in 1 ml RNAlater (Qiagen, Netherlands), and moved to a −20 °C freezer for up to a month until RNA was extracted. Left and right eyes were pooled for each individual. The pooled eyes were homogenized in a Retsch mm 400 Mixer Mill (Haan, Germany) using a carbide bead. Total RNA was extracted using the AurumTM Total RNA Fatty and Fibrous Tissue (BioRad®), which included a DNase I incubation step. The concentration and purity of the extracted RNA was assessed on a NanoDrop® Spectrophotometer (Thermo Scientific). Synthesis of cDNA was accomplished using the iScriptTM cDNA Synthesis Kit (Bio-Rad®), and 1000 ng of RNA was used as the input for the cDNA synthesis of each sample. The resulting cDNA was diluted 1:100 in ultra-pure water for RT-qPCR analysis.

To develop unique qPCR primers and probes (see Supplementary Table 1 for sequences), each opsin of the nine cone opsin genes (LWS-1, LWS-2, LWS-3, LWSr, RH2a, RH2b, SWS1, SWS2a and SWS2b) was sequenced using primers developed by (Sandkam et al. 2015). Based on these sequencing results, we designed probe and primer sets for RT-qPCR. For each gene, one of the primers and/or the RT-qPCR probe spanned an intron, which allowed us to avoid amplification of genomic DNA. We used the PrimeTime® qPCR 5’ Nuclease Assays from Integrated DNA Technologies® (Iowa, USA) for each of the targeted genes. The assays used had a double-quenched probe with 5’ 6-FAM™ dye, internal ZEN™ and 3’ Iowa Black® FQ Quencher. Using our custom primers and probes, we measured the expression of visual opsins in female fish using a standard reverse-transcription quantitative polymerase chain reaction protocol (see Supplementary Methods for full details).

Each gene’s expression was normalized against the total cone opsin expression such that each gene’s expression was represented as a percentage of the total cone opsins (see Supplementary Methods for all equations used in estimation of expression). Differences in mean expression of each opsin gene between sulphidic and non-sulphidic populations were determined using linear mixed effects models with habitat type (sulphidic or non-sulphidic) as a fixed effect and drainage as a random effect (Pinheiro et al. 2013; Ben-Shachar et al. 2020). We calculated the percent variance explained by the fixed effect using MuMIn (Barton 2009; Nakagawa and Schielzeth 2013)Since proportional cone expression is sum-constrained, we also *ln*-ratio transformed our values and repeated the linear mixed effect models (Kucera and Malmgren 1998; Veen et al. 2017). We found that results from transformed and non-transformed datasets were quantitatively similar, and non-transformed are easier to interpret, so we present figures using proportions. In addition to directly testing expression levels, we also translated opsin expression proportions into a spectral sensitivity measure using an assumption that opsins contribute additively. For each opsin, *o*, we calculated a spectral sensitivity curve *S_o_* (350-700 nm) using the absorbance templates from (Govardovskii et al. 2000) and estimates by (Kawamura et al. 2016) of the wavelength of maximum absorbance. Opsin proteins can be conjugated to the chromophores A_1_ or A_2_, which affect the shape of the absorption curve. We assume all opsins were conjugated to A_1_, rather than A_2_; we believe this is reasonable, as microspectrophotemetry by (Archer and Lythgoe 1990) found that the absorption profile of *Poecilia* visual pigments best fit the A_1_ chromophore template. These absorbance curves were summed in proportion to each opsin’s relative expression to get an individual spectral sensitivity curve for each fish.

All statistical analyses were conducted in R using tidyverse (v1.3.0) and nlme (v3.1-137) packages (Pinheiro et al. 2013; Wickham et al. 2019).

### Estimation of body colouration

As with irradiance, we smoothed reflectance measures using a rolling mean with a five nm window width and fitting a spline function to the reflectance curve from 350 to 700 nm. We removed any replicate recording with negative reflectance values. Reflectance measures were normalized so that the sum reflectance across the measured spectrum was equal across all samples. Three replicate measurements of the same region were averaged by wavelength to get a single spectrum measurement for each region on each fish. To visualize how sulphidic and non-sulphidic populations differed in coloration, we calculated the mean and standard error for reflectance at each five nm wavelength window for each population.

Reflectance across the visual spectrum is a complex phenotype with a non-independent measurement per wavelength per sample. In other systems, this type of data has been represented by the relative activation of three or four visual receptors, thereby turning a visual spectrum into a predicted perceived colour. In our case, the visual system is much more complex, because *P. mexicana* has nine visual opsins. Instead of making assumptions about colour perception, we took an agnostic approach and used a principal component analysis to describe the major axis of variation in reflectance. Reflectance measures were averaged in five nm windows (a total of 70 wavelength segments), and the principal component analysis was conducted independently for each body part including all populations together. When plotted, we found that principal component one (PC1) separated samples by environment. To test this, we conducted an ANOVA on PC1, testing for the sequential effects of drainage, habitat type, and sex, separately for each body part.

### Parallelism of opsin expression and body colouration

To determine to what degree changes in body colouration and visual sensitivity are parallel across independent drainages, we used PCA to reduce the dimensionality of the data. For body colouration, we used the mean reflectance in five nm windows as the trait values for the PCA, as described above. For visual sensitivity, we used relative opsin expression as the trait. We then used the resulting principal components (nine for opsins, 70 for colour) as input for a multivariate vector-based analysis that describes the direction of divergence between pairs of populations (Bolnick et al. 2018). In this analysis, each vector represents the direction of divergence in colour or opsin expression between the sulphidic and non-sulphidic ecotypes. A small angle between the divergence vectors of two independent ecotype pairs represents a high degree of parallelism in divergence. A 90° angle would indicate no parallelism in the pattern of divergence, and a large angle (closer to 180°) indicates an opposing direction of divergence. This vector-based approach has previously been used to estimate parallelism in phenotypes or genotypes between populations diverging repeatedly across similar environments [e.g., (Stuart et al. 2017; Rennison et al. 2019)]. We described the direction of divergence between sulphidic and non- sulphidic ecotypes within each drainage using a vector connecting the mean position (centroid) of individuals of one ecotype to the mean position of individuals of the other ecotype. We estimated the angle (θ, in degrees) between the divergence vectors of each ecotype pair (from each drainage) and calculated the average angle. To assess parallelism, we then tested whether the average angle between divergence vectors of different ecotype pairs was smaller than expected by chance. We used t-tests to determine significance and 90° as the null or random expectation.

### Correlation between visual sensitivity, light absorbance, and body colouration

We found differences in visual sensitivity, light absorbance, and body colouration between sulphidic and non-sulphidic populations, so we next asked if these shifts were correlated between different spectral quantities of the water bodies. For example, is decreased short wavelength sensitivity in sulphidic populations accompanied by decreased short wavelength light availability? We answered this question by taking a spectrum wide approach described fully in the supplementary material of Rennison et al. (2016) and diagrammed in Supplementary Figure 1. We used a statistic to quantify the association between the shift in spectral sensitivity, changes to body reflectance, and the transition in light environment between sulphidic and non-sulphidic populations across all wavelengths for each drainage. For each population, we constructed transmission (K_s_) curves by calculating at each wavelength (λ) the median K_s_ for all samples within a location. At each wavelength, we then subtracted the median value of the sulphidic location from the median value of the non-sulphidic location within a drainage, yielding the change in transmission (ΔK_s_). Change in spectral sensitivity (ΔS) was calculated similarly as follows. For each population, we calculated the median sensitivity at each wavelength using the proportions of opsin expression and maximal sensitivity assuming an A1 chromophore. Change in sensitivity was calculated as the difference between the median non-sulphidic and sulphidic sensitivity curves. Lastly, for the change in body colour or reflectance (ΔR), we calculated the median reflectance per wavelength per population per body part and again subtracted the non-sulphidic population from the sulphidic population values.

This resulted in three spectral quantities—transmission, sensitivity and reflectance—measuring the difference between sulphidic and non-sulphidic populations in each drainage. For reflectance, we have four different measures for the four body parts recorded. We chose pairs of spectral quantities and calculated the correlation coefficient (*r*) between them. For example, a positive *r* indicates that regions of the spectrum with increased transmission also have increased reflectance or sensitivity. We tested if *r* was significantly different from zero (no relationship) for each combination, using drainage as our unit of replication.

## Results

### Opsin expression and visual sensitivity

In all samples, opsin expression was predominantly violet sensitive SWS1, blue sensitive SWS2B, green sensitive RH2-1, and green sensitive LWS-3 (Supplementary Figure 2). We compared proportional expression of opsins between sulphidic and non-sulphidic environments, while controlling for drainage, and found significant (P < 0.05) differences between populations from different habitat types in RH2-1 and LWS-3, and a strong trend of differences in SWS2B expression (Figure 2A-C, Table 1). For these three genes, the direction of divergence in opsin expression was repeated across the four independent drainages.

**Table 1.** Difference in mean percent cone opsin expression between sulphur and non-sulphur populations. Results of linear mixed effect models using proportional cone expression. P-values included for *ln*-ratio transformed cone expression are quantitatively similar to non-transformed.

| Gene | $\lambda_{\text{max}}$<br>(nm) | Difference in<br>expression (+/-<br>STE) | $F_{1,51}$ | Effect size<br>95% C.I. | Percent<br>variance<br>explained | p (transformed p) |
| --- | --- | --- | --- | --- | --- | --- |
| LWS-1 | 571 | 1.78E-5 (4E-5) | 0.20 | [-0.21, 0.33] | <0.01 | 0.65 (0.47) |
| LWS-2 | 516 | 7.9E-5 (2E-4) | 0.25 | [-0.33, 0.20] | <0.01 | 0.62 (0.32) |
| LWS-3 | 519 | 0.05 (0.01) | 50.45 | **[-0.80, -0.45]** | **0.37** | **< 0.0001 (< 0.0001)** |
| LWSr<br>(LWS-4) | NA | 8.9E-6 (6E-5) | 0.02 | [-0.28, 0.24] | <0.01 | 0.89 (0.16) |
| RH2-1 | 516 | 0.14 (0.03) | 25.03 | **[0.29, 0.68]** | **0.53** | **< 0.0001 (<0.0001)** |
| RH2-2 | 476 | 0.002 (0.005) | 0.19 | [-0.20, 0.31] | <0.01 | 0.66 (0.08) |
| SWS1 | 438 | 0.01 (0.02) | 0.71 | [-0.36, 0.15] | 0.01 | 0.41 (0.12) |
| SWS2A | 408 | 0.001 (0.001) | 1.04 | [-0.39, 0.13] | 0.02 | 0.31 (0.53) |
| SWS2B | 353 | 0.08 (0.04) | 3.72 | [-0.52, 0.01] | 0.06 | 0.06 (0.29) |

**Figure 2:**
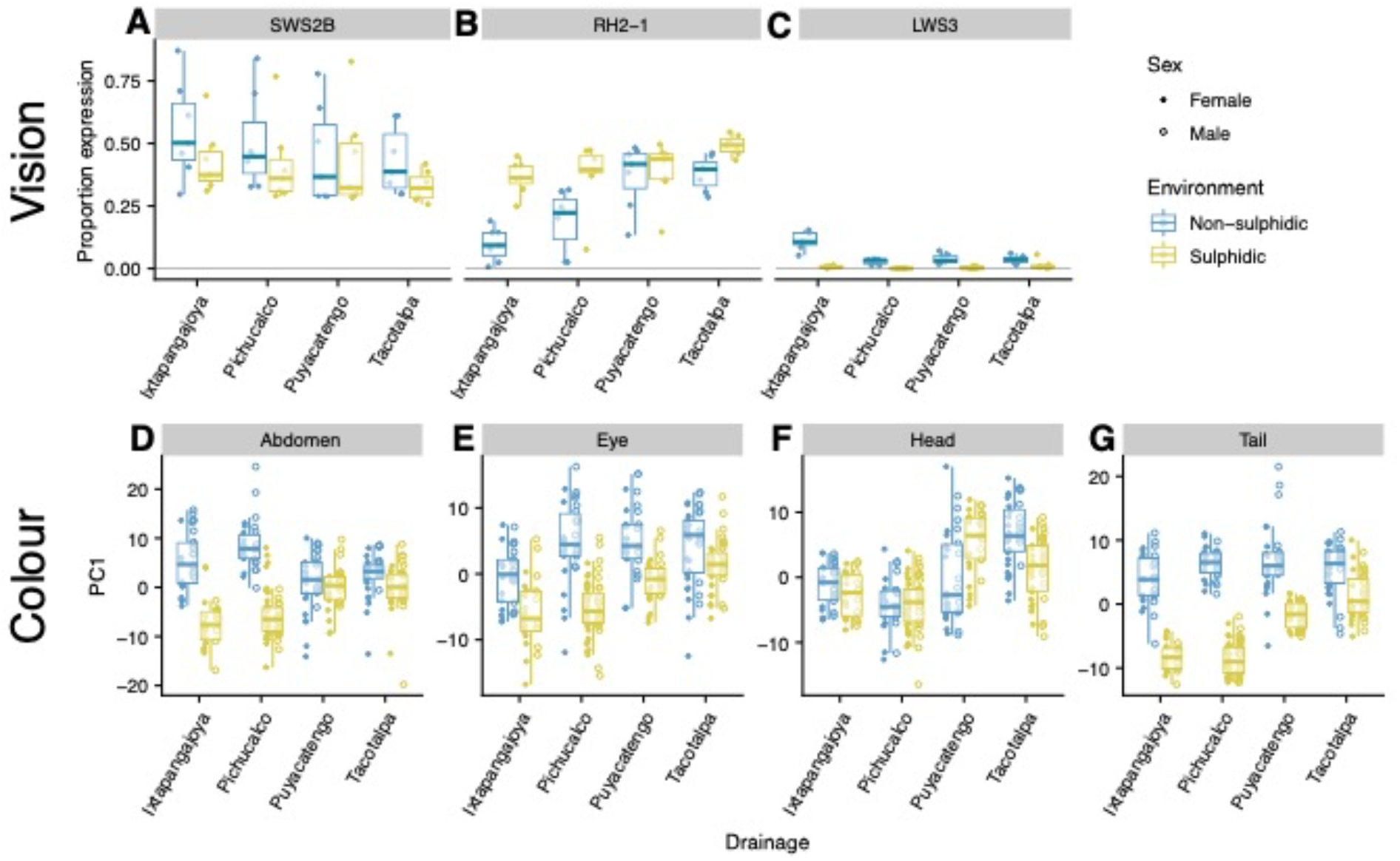
Parallel phenotypic differentiation in vision and body colour between environments. A-C: Proportion opsin gene expression for genes differentially expressed between environments. D-G: Principal component 1 scores for body colour. Box area contains the middle two quantiles.

Based on opsin expression, we calculated sensitivity curves for all samples (Supplementary Figure 3). Inferred sensitivity peaked at 438 and 516 nm, corresponding to the three highly expressed genes. Due to the consistent differences in opsin expression, we found generally more long-wavelength sensitivity in sulphidic populations and more short-wavelength sensitivity in non-sulphidic populations.

### Body colouration

Principal component analysis was effective at reducing dimensionality of our reflectance spectrum measures. For each body part, the first two principal components (PCs) explained between 85.4% and 92.9% of the total variation (Table 2; Supplementary Figure 4). In most cases, the first principal component, which explained more than half of the variation, separated samples by habitat type (sulphidic vs. non-sulphidic) (Figure 2D-H). We further probed the sequential effect of drainage, habitat, and sex using an ANOVA of PC1 and found that the environment explained the most variation in abdomen, eye, and tail colouration (Table 2). In all cases, sex played a relatively small role in explaining variation of the first PC, although we note that our measurements don’t include the dorsal or caudal fins, where orange/yellow pigmentation is sexually dimorphic. From examining the spectrum, we found that for body parts that differed based on environment (the abdomen, eye, and tail), there was generally more reflectance of short wavelengths in non-sulphidic environments than sulphidic ones, and the opposite pattern for long wavelengths (Supplementary Figure 5).

**Table 2:** The analysis of variance in PC1 of body colouration.

| *Body Part* | Drainage |  |  | Sulphide |  |  | Sex |  |  |
| --- | --- | --- | --- | --- | --- | --- | --- | --- | --- |
|  | *F-value* | *p* | *PVE* | *F-value* | *p* | *PVE* | *F-value* | *p* | *PVE* |
| Abdomen | 2.97 | **3.21E-02** | 1.99 | 142.42 | **8.74E-27** | 31.79 | 18.66 | **2.18E-05** | 4.16 |
| Eye | 23.96 | **8.01E-14** | 14.82 | 114.57 | **1.29E-22** | 23.62 | 20.53 | **8.74E-06** | 4.23 |
| Head | 46.84 | **1.63E-24** | 32.88 | 1.24 | 2.66E-01 | 0.29 | 8.58 | **3.69E-03** | 2.01 |
| Tail | 56.45 | **1.59E-28** | 19.07 | 434.17 | **9.99E-59** | 48.89 | 6.49 | **1.14E-02** | 0.73 |

### Light environment

Irradiance measures indicated that the ambient light in both environments was broad spectrum and spanned from ultraviolet to the long wavelengths of the visual spectrum (Supplementary Figure 6). Based on repeated measures at depths up to 50 cm, we found nearly perfect light transmission; light absorbance by water bodies was minimal, and any absorption was limited to short wavelengths (350 to 400 nm) (Supplementary Figure 7). This means that there was no detectable loss of light due to depth for wavelengths > 400 nm, which resulted in practically no difference in the transmission profiles of sulphidic and non-sulphidic habitats within each drainage for most of the light spectrum. For short wavelength light, transmission was higher in sulphidic water, which would lead to greater short wavelength light availability.

### Parallel phenotypic change

We used a vector-based analysis of PCA space to quantify the degree of parallelism between pairs of populations for opsin expression and body colouration (Figure 3A). We combined all of the opsin expression data into a PCA analysis and used the resulting loadings to quantify the degree of parallelism in the overall direction of divergence between sulphidic and non-sulphidic population pairs. We found significant parallelism across the four independent replicates (mean θ = 70.5°, range: 62.6° - 83.1°, p = 0.002, t_5_ = −5.95; Figure 3B). There was also significant parallelism in the direction of divergence in body colouration between replicate sulphidic and non-sulphidic populations for the tail (mean θ: 35.5°, range 8.5° - 57.0°, p = 0.002, t_5_ = −6.06), abdomen (mean θ: 43.0°, range: 16.0° - 72.7°, p < 0.0001, t_5_ = −12.33), and eye (mean θ: 40.0°, range: 28.1° - 53.0°, p = 0.002, t_5_ = −5.95). There was no evidence for parallelism in the direction of divergence for the head colouration (mean θ: 85.38°, range: 48.8° - 132.3°, p = 0.71, t_5_ = −0.39; Figure 3C-F). The degree of parallelism tended to be greater for body reflectance than for opsin expression. The magnitude of parallelism also varied among pairwise population comparisons for each trait. In general, parallelism tended to be greater for population pair comparisons that did not include the Tacotalpa drainage (Figure 3). The results of the vector-based parallelism analysis were consistent when male and female pigmentation was analysed separately (results not shown).

**Figure 3:**
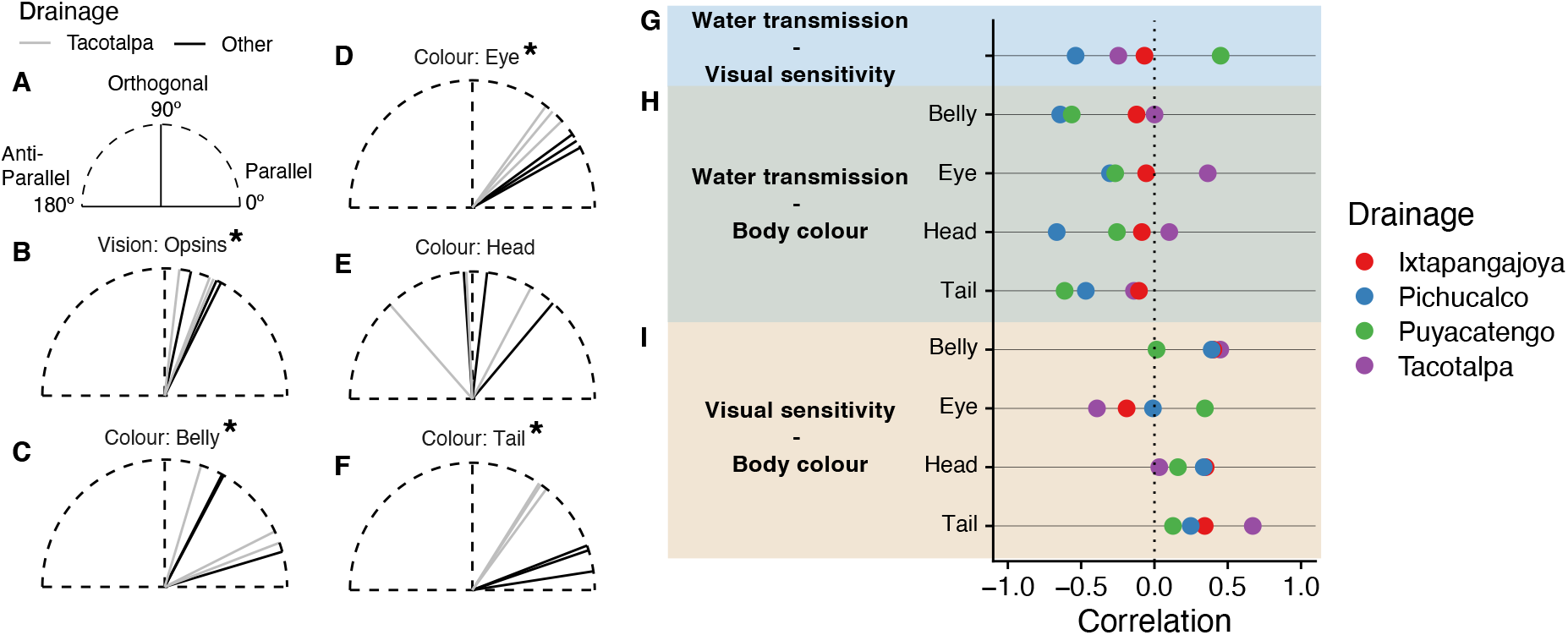
Parallel trait evolution and correlation between spectral quantities. A: Evolutionary trajectories diagram. B-F: Pairwise evolutionary trajectories for vision and body colour phenotypes. Each line represents a pair of drainages. * indicate trajectories significantly different from 90° (p < 0.003). Comparisons including the Tacotalpa drainage are indicated by grey colouration. G-I: Correlation between different spectral quantities.

### Correlations between visual sensitivity, light availability and body colouration

We found that water transmission and either visual sensitivity or body colour (i.e., reflectance) were generally negatively correlated (16 out of 20 comparisons) (Figure 3G,H). This result indicates that—for a given wavelength—reduced light availability tended to be associated with increased visual sensitivity or pigmentation reflectance. In contrast, visual sensitivity and body colouration were generally positively correlated (13 out of 16) (Figure 3I), indicating that increased visual sensitivity tended to be associated with increased reflectance of body pigmentation. Out of our four drainages, Tacotalpa most often had a correlation pattern counter to the other three drainages; i.e., negative where others are positive. We tested each comparison to see if the mean correlation was different than zero using a two-sided student t-test and found that four of the comparisons were near significant (0.05 < p < 0.09) (Supplementary Table 2), including three of the comparisons between visual sensitivity and body colouration.

## Discussion

### Parallel phenotypic shifts

Repeated shifts in the phenotypes and/or genotypes of organisms that have independently colonized new environments provides strong evidence for the action of natural selection and suggests adaptive value (Schluter and Nagel 1995). In our survey of four independent drainages containing sulphidic and non-sulphidic populations of molly, we found consistent differences in the opsin gene expression levels and body colour reflectance between the two ecotypes. We see repeated differences in the expression of the SWS2B (λ max 353 nm) and RH2-1 (λmax 516 nm) opsins, as well as the LWS-3 (λmax 519 nm). Differences between the ecotypes in the expression of the other six opsins appear to be drainage specific. Together, the differences in opsin expression between sulphidic and non-sulphidic populations are predicted to translate into differences in the overall visual sensitivity, and perhaps discriminatory ability, of the two ecotypes (Supplementary Figure 3). Functionally, the parallel shifts in visual sensitivity appear to have reduced short wavelength sensitivity, while comparatively increasing medium wavelength sensitivity of sulphidic populations. Parallel shifts in opsin expression have previously been described in several fish species, including African cichlids (O’Quin et al. 2010), threespine stickleback (Rennison et al. 2016), and Neotropical Midas cichlids (Torres-Dowdall et al. 2017). This suggests that the forces shaping opsin expression (and correspondingly spectral sensitivity) are often fairly consistent across habitat transitions. Shifts in body colour reflectance followed a similar pattern, with sulphidic populations reducing short-wavelength reflectance while increasing medium- and long-wavelength reflectance of patches behind the eye, the tail, and abdomen. Differences in body pigmentation are often seen between organisms inhabiting different niches or habitats and are considered a classic example of parallel evolution (Hoekstra 2006).

Observed shifts in pigmentation and opsin expression between sulphur and non-sulphur molly ecotypes could be due to genetic and/or plastic changes. Previous work has in fish has shown that variation in opsin expression between fish occupying different light regimes can be largely heritable (e.g. threespine stickleback (Flamarique et al. 2013; Rennison et al. 2016), damselfish (Stieb et al. 2016); Atlantic molly (Tobler et al. 2010)). or largely plastic (e.g. African cichlids (Nandamuri et al. 2017); red shiner (Chang and Yan 2019); cardinalfish (Luehrmann et al. 2018)). Plasticity seems key for responses to short term or small-scale variation in light environment (e.g. (Stieb et al. 2016; Veen et al. 2017; Kranz et al. 2018)). Experimental work in guppies suggests that heritable change in opsin expression may require several generations of exposure to a differential light environment (Kranz et al. 2018). It is likely that the differences in opsin expression we observe between sulphur and non-sulphur ecotypes is likely due to a combination of genetically determined factors and phenotypic plasticity, although we found only minimal differences in light spectrum between environments, suggesting that short-term plastic responses to light environments are unlikely to explain the parallel shifts. Future efforts should work toward quantifying the relative contribution of heritable and non-heritable change.

The angle (magnitude) of parallelism was highly variable across traits and among the four drainages. A small angle between two divergence vectors indicates a very similar pattern of divergence between two independent ecotype pairs. We found that four of the five traits surveyed diverged in a significantly parallel manner across the independent drainages. However, the patterns of divergence across drainages were most similar for tail, abdomen, and eye reflectance (average angles of divergence were 35°, 43° and 40°, respectively). The pattern of divergence in opsin expression, while significantly parallel, was considerably less parallel than that seen for the three pigmentation traits with an average angle of 70.5° between pairwise vectors. This suggests that the selective forces shaping patterns of differentiation in body pigmentation are perhaps more consistent among drainages than those affecting opsin gene expression. The vector-based approach used here has been previously applied in threespine stickleback to quantify the pattern of parallel evolution of morphological traits. For comparison, quantification of morphological parallelism (based on 84 phenotypic traits) across 16 replicate stream and lake ecotype pairs of threespine stickleback revealed multivariate angles ranging from 30° to 135° between any two ecotype pairs (Stuart et al. 2017); thus, it appears the parallelism of pigmentation in this system is fairly strong relative to that described for other phenotypes.

The similarity of the selective landscape appears to be variable with certain drainages exhibiting a more unique pattern of divergence (or lack of divergence) than the others. Within a trait, there was often considerable difference in the angles of pairwise divergence vectors. For example, the angle between pairs of vectors describing divergence in tail reflectance ranged from 8.5° to 57° and from 48° (parallel) to 132° (antiparallel) for head reflectance. Interestingly, across the traits, comparisons involving the Tacotalpa drainage tended to have larger angles than those based on the other three drainages. This may in part be explained by the fact that the Tacotalpa drainage does not only contain non-sulphidic and sulphidic ecotypes, but *P. mexicana* have also colonised and adapted to a non-sulphidic and a sulphidic cave (Tobler et al. 2008). Cave populations are characterised by regressive evolution of body pigmentation and eye function, including reduced opsin gene expression (Tobler et al. 2010; McGowan et al. 2019), and are connected to the sulphidic surface population investigated here by low levels of gene flow (Tobler et al. 2008). Hence, introgression of alleles from populations exhibiting different selective environments (i.e., the absence of light) might contribute to the unique evolutionary trajectory of the Tacotalpa population.

### Correlated environmental and phenotypic shifts

Given our finding of parallel shifts in pigmentation and opsin expression, we sought to determine whether the two traits were co-evolving and whether shifts in these traits corresponded with differences in the spectral environments of sulphidic and non-sulphidic habitats. Across the four drainages, there were positive correlations between shifts in visual sensitivity and shifts in body pigmentation This pattern was found for abdomen, eye, and tail reflectance (although none were statistically significant), suggesting that these three pigmentation traits and spectral sensitivity may be co-evolving. Divergence in tail and abdomen colouration also tended to correlate with differences in light environment, with the four drainages showing a similar pattern. For the other two traits, eye and head colour, there were consistent correlations with differences in ambient light for three of the four drainages. Interestingly, the pattern of divergence for these traits was mismatched (positively correlated) for only the Tacotalpa drainage, where we also documented less parallelism. Overall, our findings suggest that differences in the properties of ambient light between sulphidic and non-sulphidic environments may contribute to patterns of divergence in pigmentation, although we are limited by the number of replicate populations. The generally negative correlations imply that greater light absorbance, which results in less light availability, is correlated with greater sensitivity or greater reflectance at those wavelengths. Further testing will be required to determine whether a function, such as crypsis, relates to the match with ambient light. In many examples of repeated pigmentation changes, crypsis is evoked as the driver of such shifts. For example, several mouse and lizard species exhibit loss of pigmentation after colonization of light sand environments from darker substrates (reviewed by Hoekstra, 2006).

Matching of differences in ambient light conditions and spectral divergence and/or body pigmentation may be further evidence of local adaptation of these traits. A match between spectral sensitivity and available wavelengths has long been predicted, as it is thought to maximize an individual’s photon catch (Clarke 1957; Denton and Warren 1957; Munz 1958). However, evidence of this matching has been limited. The first comprehensive test of the hypothesis using spectrum wide data, rather than just maximal absorbance values, found moderate spectral matching in threespine stickleback (mean *r* = 0.39) (Rennison et al. 2016). In mollies, we find little support (mean *r* = −0.05) for spectral matching. Correlations between shifts in visual sensitivity and differences in the light transmission between sulphidic and non-sulphidic habitats were negative for three of four drainages (*r* = −0.26 to 0.16). This indicates that in regions of the spectrum where there was decreased transmission, there was increased visual sensitivity, perhaps to compensate. Variable and weak matching of spectral sensitivity on ambient light parameters suggest that other visual functions, such as contrast detection and colour discrimination, are also likely to be important in shaping spectral sensitivity (Loew and Lythgoe 1978). However, a drawback of the study is that sampling of ambient light was conducted at one time point and seasonal fluctuations could play an important role in determining patterns of divergence. We measured ambient light during the dry season when there is low flow and very low turbidity. During the wet season, turbidity increases with flow and more dramatic visual changes to water clarity are present; sulphidic waters usually acquire a blue, milky turbidity, while non-sulphidic waters shift to warmer earth-tones (Figure 1). Difference in light transmission during this time may be different from our current estimates and may help to explain the pattern of divergence.

Sensory drive and sensory bias models have been used to explain correlated patterns of divergence of sexual signals and sensory systems. These models predict positive correlations between female perception, male sexual signals, and the signalling environment (in the case of sensory drive) (Boughman 2001). Here, we found that divergence of body reflectance in several body regions are indeed accompanied by matched shifts in spectral sensitivity of female fish and correlate with parameters that describe the signalling environment. However, sensory drive and sensory bias models often consider sexually dimorphic traits (e.g., (Boughman 2002; Seehausen et al. 2008)). Curiously, we find that male *and* female fish exhibit similar phenotypic patterns for the pigmentation traits included in our study and correspondingly have similar patterns of trait divergence and matching. Molly populations often exhibit sexual dimorphism in pigmentation (Figure 1). One possible reason why we did not find sexual dimorphism in pigmentation is that male nuptial colours are flexible and can be lost between capture and measurement due to stress. Additionally, sexually dimorphic pigmentation patterns may primarily be on the dorsal and caudal fins which were not measured in this study due to measurement issues with background reflectance through transparent fin tissues. Nevertheless, it is possible that females exhibit preferences towards certain pigment patterns, as female preference evolution has been seen in other contexts, which leaves the possibility that sexual selection contributes to divergence of these traits between sulphidic and non-sulphidic populations (Plath et al. 2006). Further experimental work will be required to explicitly test whether there is evidence for variation in female preference for pigmentation traits. Such tests will be pivotal in evaluating whether this system exemplifies sensory drive or sensory bias.

Ecological processes aside from sensory drive/bias may also explain the putatively adaptive shifts in both visual capacity and pigmentation. Sulphidic and non-sulphidic habitats differ in their food webs, fish communities, and levels of bird predation (Riesch et al. 2010a; Tobler et al. 2015). Different predator communities could affect overall predation risk and correspondingly the need for crypsis. Previous work has documented the evolution of behavioural changes in response to these changed predation pressures (Lukas et al. 2021). Further, the differential absorption of short wavelengths of light in sulphidic habitats may affect what pigmentations best avoid detection by predators. Differential nutrient acquisition between habitats could also affect the expression of pigmentation between sulphidic and non-sulphidic populations, as has been suggested for guppies (Grether et al. 2001). Experimental work isolating these different agents of selection, which are correlated in nature, will be required to determine the most proximate mechanisms underlying our observed patterns.

## Conclusions

We surveyed the divergence of spectral sensitivity and body pigmentation for four replicate population pairs of mollies inhabiting sulphidic and non-sulphidic habitats. We find robust evidence of parallel shifts in opsin gene expression, predicted spectral sensitivity and body reflectance. Shifts in vision seem to be accompanied by correlated shifts in pigmentation, which are also somewhat matched to features of ambient light. Thus, we suggest that these differences are likely adaptive. Further work will be required to determine whether both natural and sexual selection contribute to the observed patterns and what specific selective agents contribute to differential fitness.

## Data availability

All data and code are available on github at https://github.com/djrennison/sulphide_molly

## Acknowledgements

We would like to acknowledge the Society for Experimental Biology for a Company of Biologists Travel grant which funded the field work conducted by DJR. This work was supported by grants from the National Science Foundation (IOS-1557860, IOS-1931657).

## Author Contributions

D.J.R., G.LO. and M.T. conceived of the project. D.J.R., and M.T. collected samples and environmental measurements. G.L.O, D.J.R., and D.R.M conducted molecular lab work. G.L.O., D.J.R. and T.V. analysed resulting data. L.A.R facilitated the field collections. G.L.O. and D.J.R. wrote the manuscript with input from all authors.

## Supplementary Material

**Supplementary Table 1.**
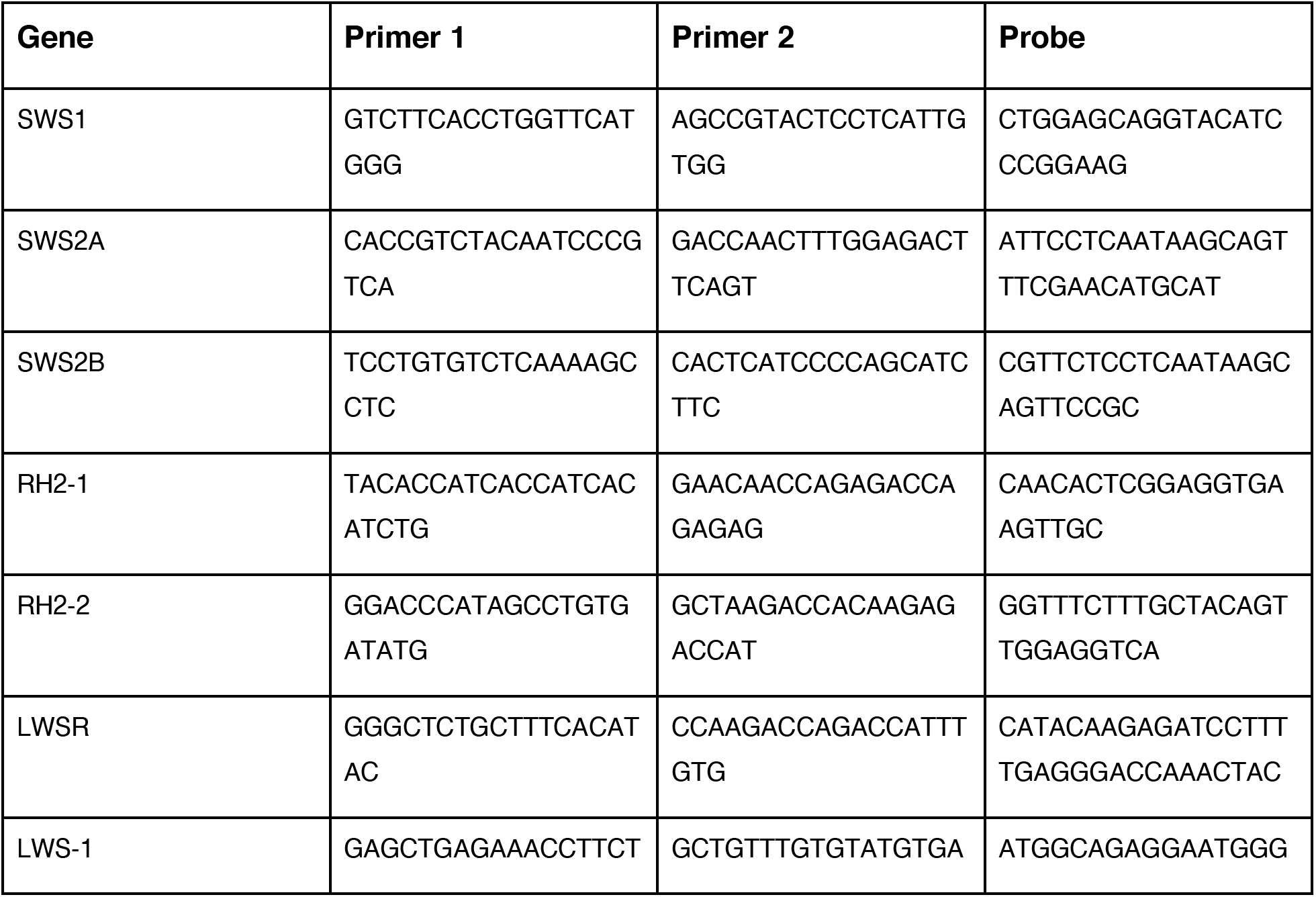
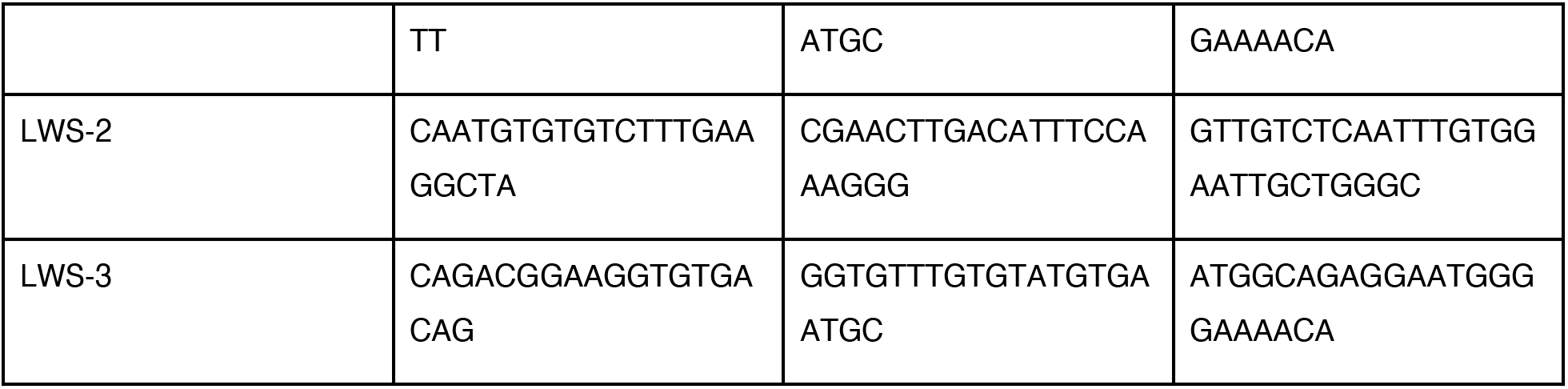
Primer and probe sequences used for the RT-qPCR assay of each opsin gene.

**Supplementary Table 2:**
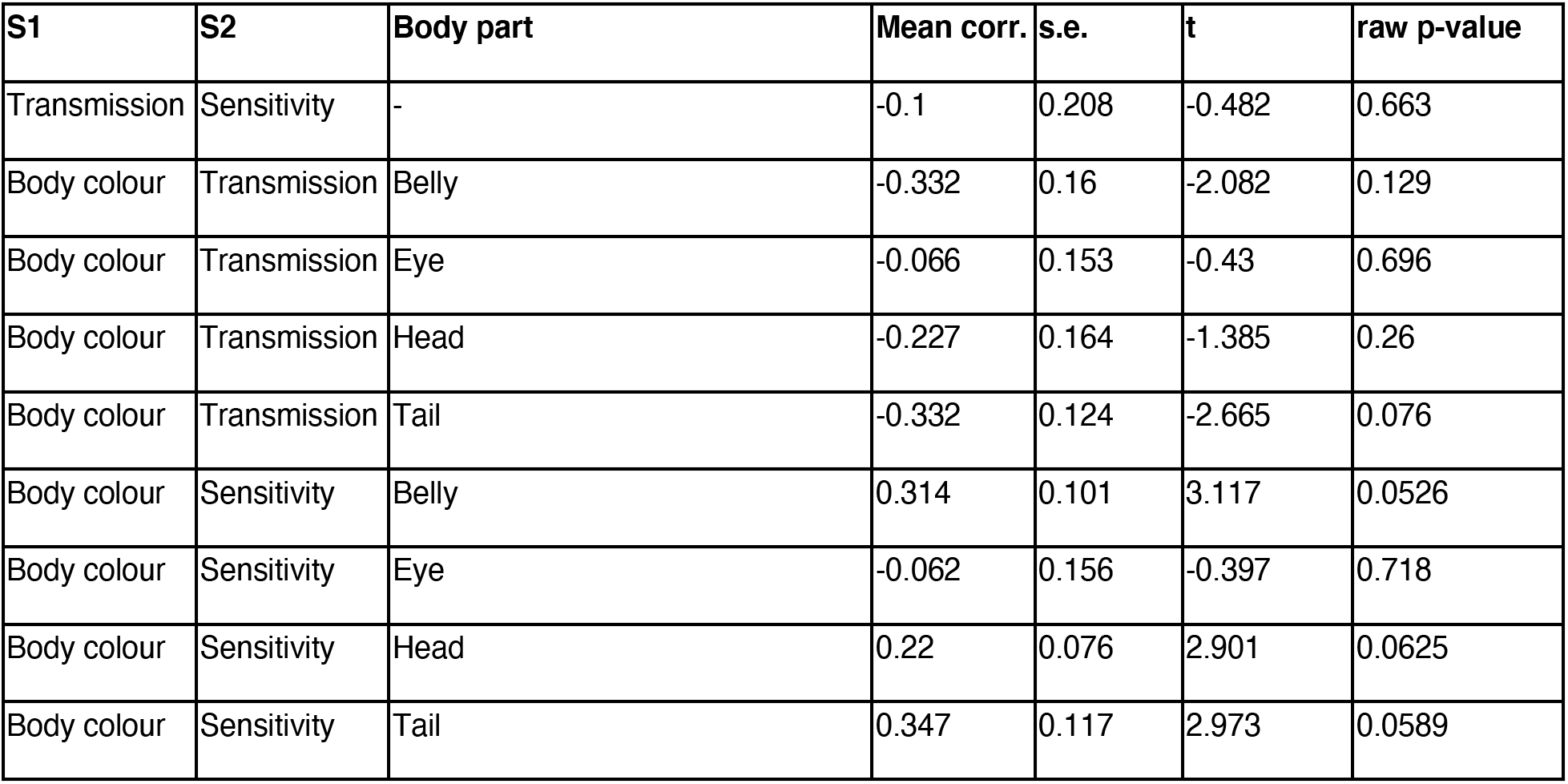
Mean correlation between the change in spectral quantities from non-sulphur to sulphur populations.

| S1 | S2 | Body part | Mean corr. | s.e. | t | raw p-value |
| --- | --- | --- | --- | --- | --- | --- |
| Transmission | Sensitivity | - | -0.1 | 0.208 | -0.482 | 0.663 |
| Body colour | Transmission | Belly | -0.332 | 0.16 | -2.082 | 0.129 |
| Body colour | Transmission | Eye | -0.066 | 0.153 | -0.43 | 0.696 |
| Body colour | Transmission | Head | -0.227 | 0.164 | -1.385 | 0.26 |
| Body colour | Transmission | Tail | -0.332 | 0.124 | -2.665 | 0.076 |
| Body colour | Sensitivity | Belly | 0.314 | 0.101 | 3.117 | 0.0526 |
| Body colour | Sensitivity | Eye | -0.062 | 0.156 | -0.397 | 0.718 |
| Body colour | Sensitivity | Head | 0.22 | 0.076 | 2.901 | 0.0625 |
| Body colour | Sensitivity | Tail | 0.347 | 0.117 | 2.973 | 0.0589 |

**Supplementary Figure 1:**
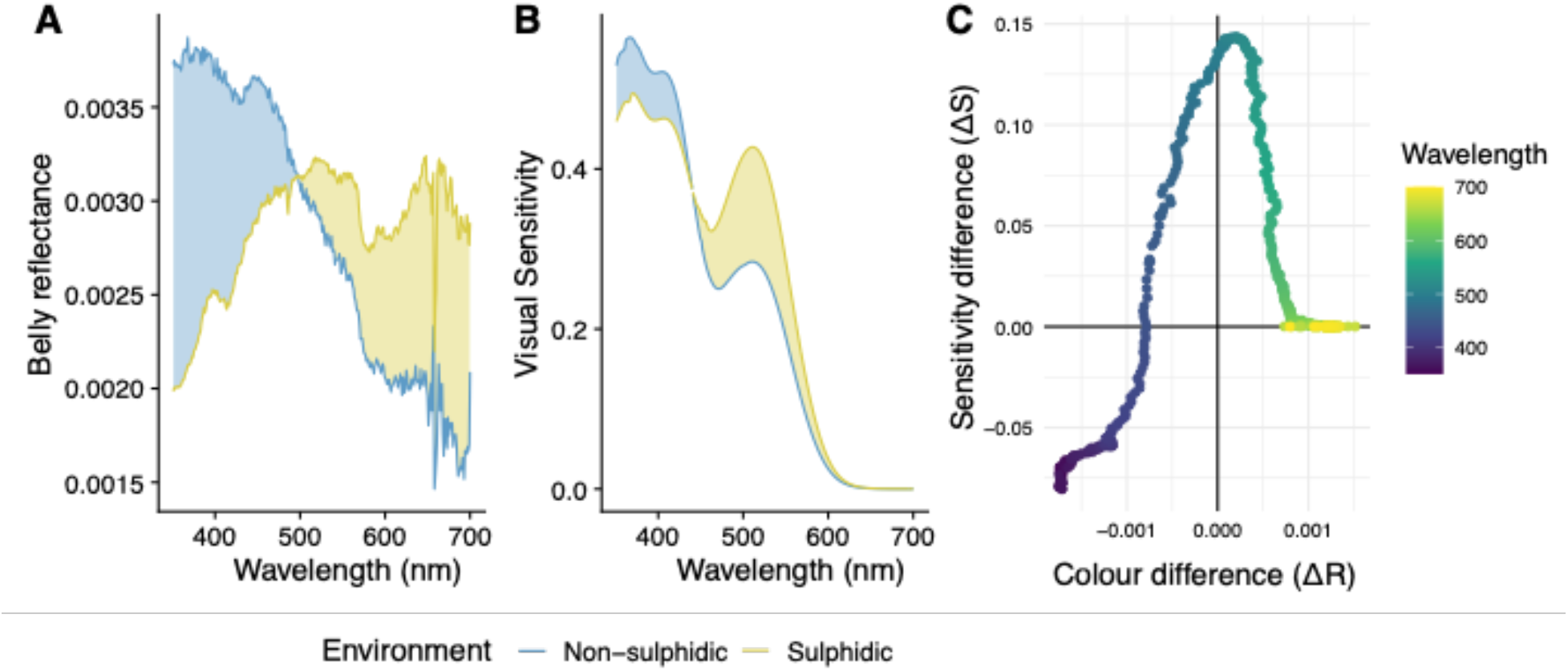
An example of a spectral correlation analysis. A) Median normalized belly colour for sulphidic and non-sulphidic populations of a single drainage. The difference between reflectance (ΔR) is highlighted. B) Median inferred visual sensitivity between sulphidic and non-sulphidic populations of a single drainage. C) The correlation between ΔS and ΔR.

**Supplementary Figure 2:**
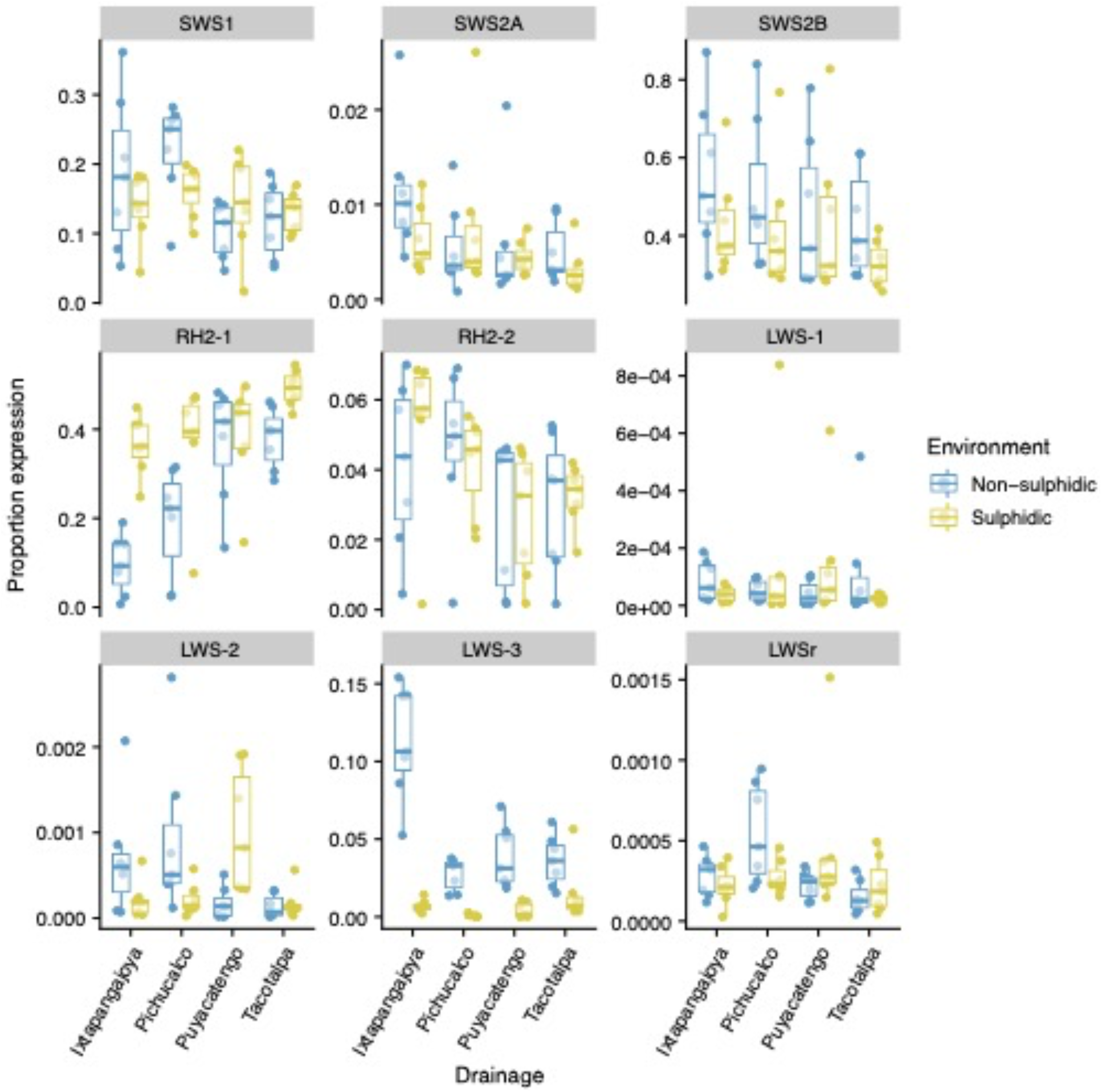
Proportional opsin gene expression in *Poecilia* samples. Opsin gene expression normalized by total opsin expression.

**Supplementary Figure 3:**
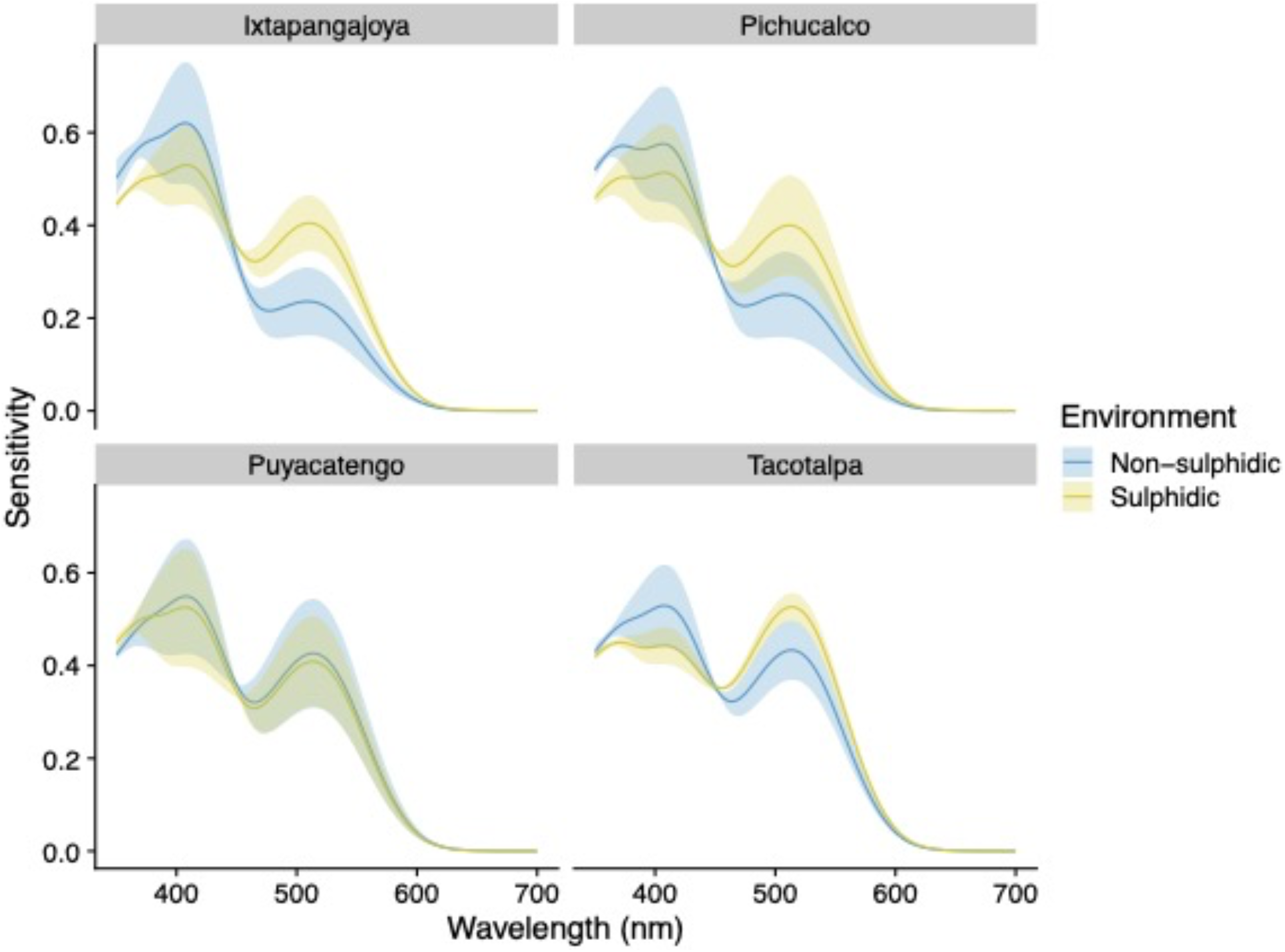
Estimated spectral sensitivity based on opsin gene expression. All opsins were assumed to be conjugated to A1 chromophore. The solid line indicates the mean estimated sensitivity for each wavelength and population and the shaded area is the 95% confidence interval.

**Supplementary Figure 4:**
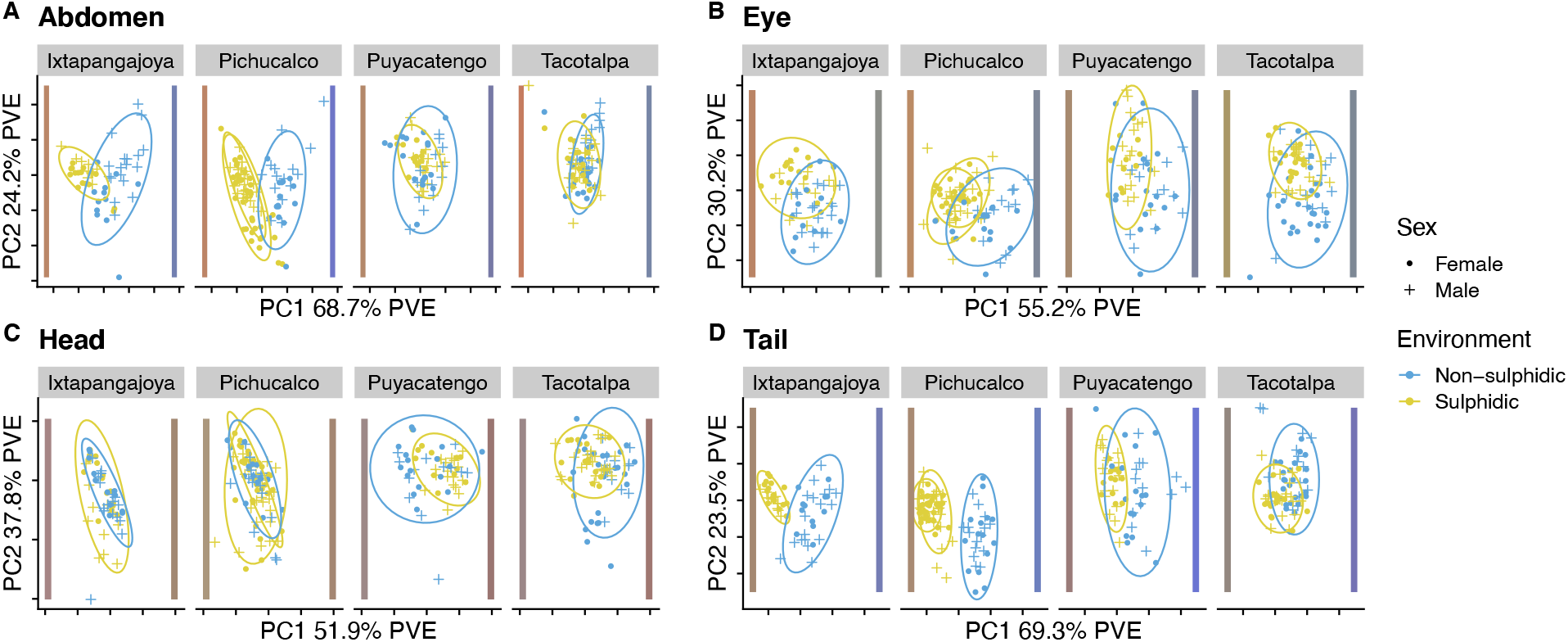
Principal component analysis of skin reflectance. Median reflectance per sample in 5 nm windows from 350 to 700 nm was used as the input data. Each body part was analysed separately. Ovals represent 95% confidence intervals and coloured bars are colour representations of the reflectance spectra of samples with the minimum and maximum PC1 scores.

**Supplementary Figure 5:**
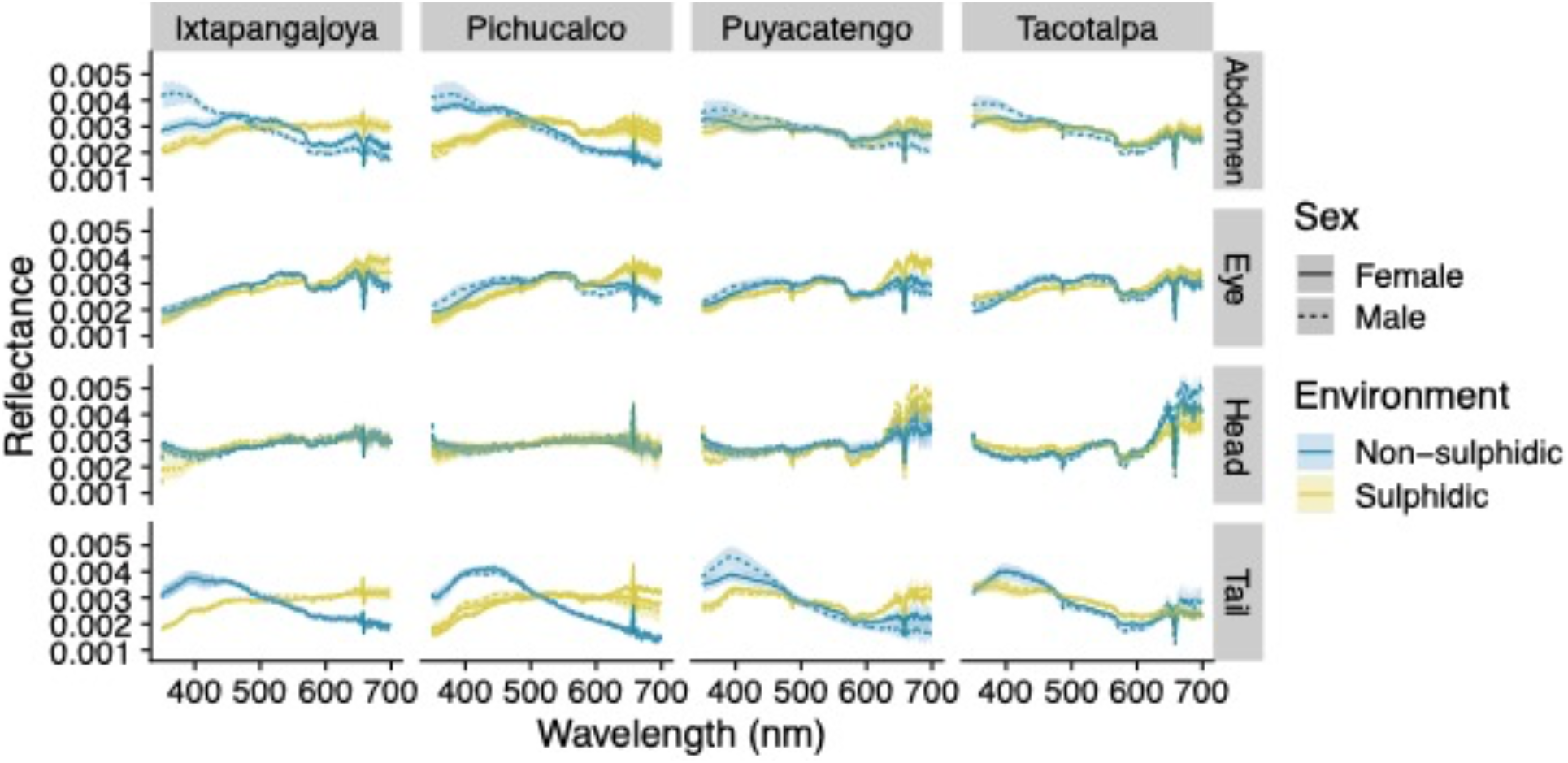
Normalized body reflectance. The line indicates the mean normalized reflectance value and the shaded value is the 95% confidence interval.

**Supplementary Figure 6:**
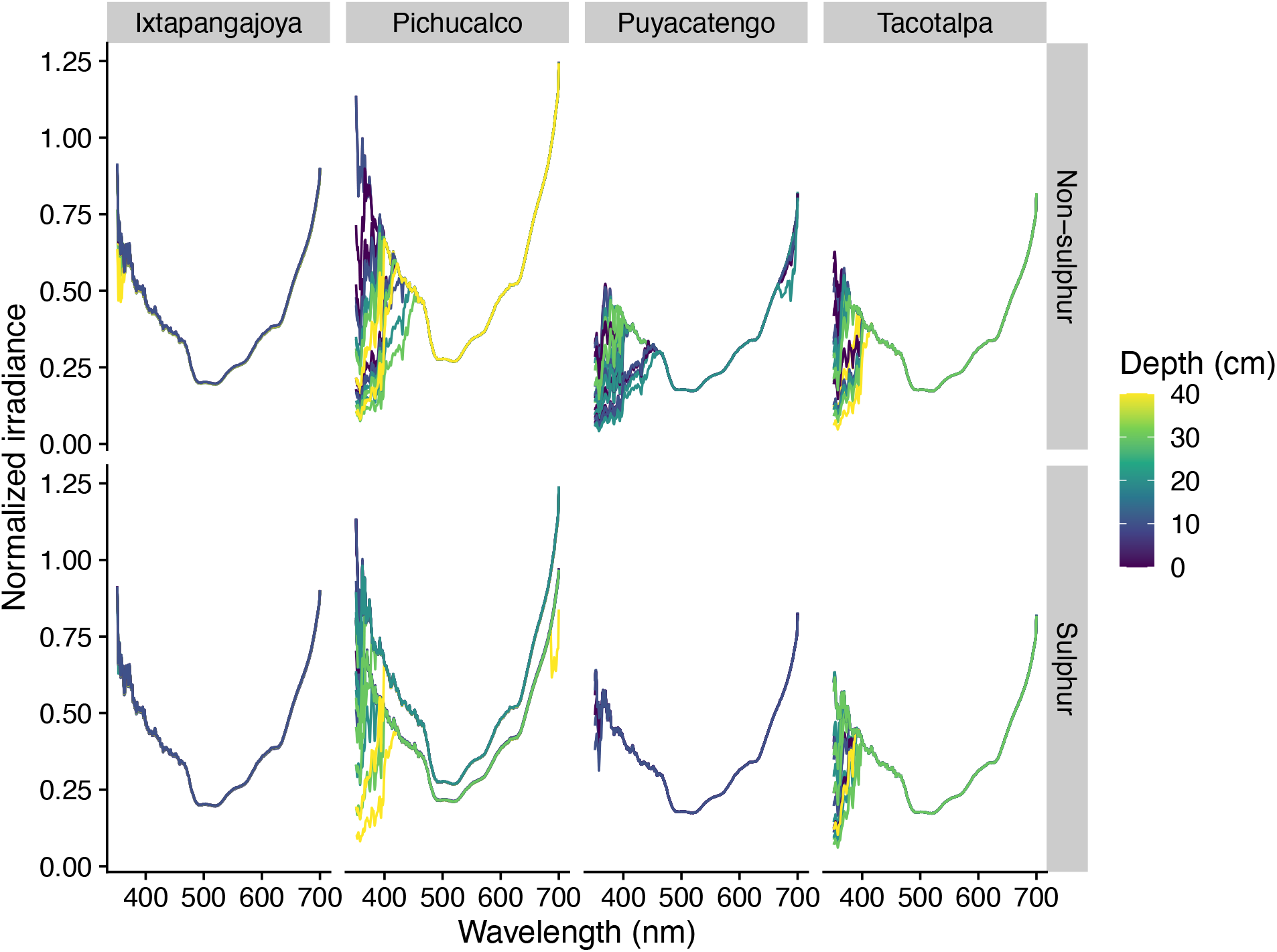
Normalized irradiance spectrum for collection sites. Each line represents the spline-fitted value for a single side-welling measurement within a location. Lines are coloured by the water depth.

**Supplementary Figure 7:**
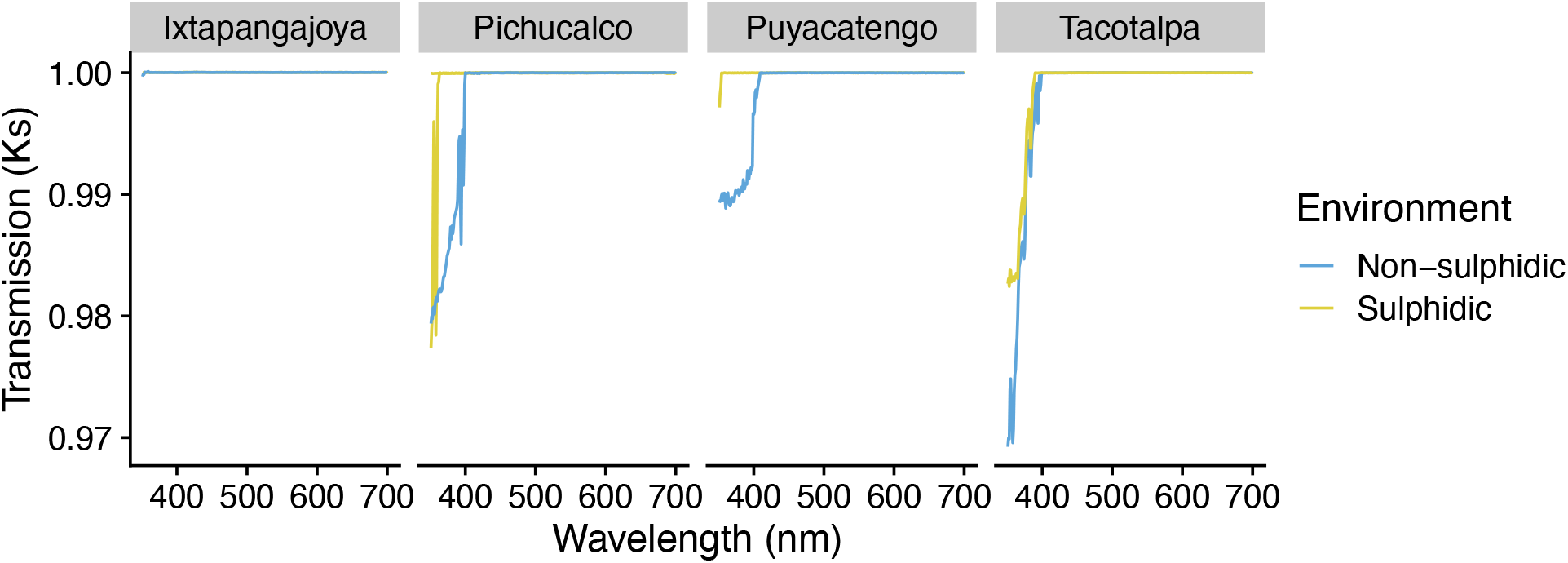
Median light absorbance for measured sites. Lower values mean more photons are lost during transmission through the water.

## Supplementary Methods

Quantification of gene transcript copy number was done using RTqPCR analysis on a BioRad®IQ5 machine (BioRad, California USA). The polymerase used was the SsoFast probes supermix (BioRad®) in a 25 μl reaction. Reactions were run in 96-well plates (Fisher, Massachusetts USA), which were sealed using optical sealing tape (BioRad®). Well-factors were collected from each of the experimental plates. Reactions were run in duplicate or triplicate. No-reverse transcription and no template controls were included for every run and did not amplify. RT-qPCR conditions consisted of 1 cycle at 95 C (3 minutes); 40 cycles of 95 C (10 seconds) followed by 60 C (30 seconds). We used a standardized luminance threshold value of 50 to calculate CT values. Equation 1 was used to calculate the PCR efficiencies (E) for each of the primer pairs,

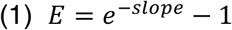

where the slope is determined from a linear least squares regression fit to critical threshold (Ct) data from a cDNA dilution series (1:10, 1:50, 1:100, 1:500, 1:1000).

For the purposes of this study we were more interested in the expression of each opsin gene relative to the total opsin levels present in the retina, rather than absolute levels of expression, so we used the proportion of total opsin expression for a given gene. The estimate of the initial amount of gene transcript (Ti) was calculated for each individual (i) using equation 2, where E is the PCR efficiency for a given gene calculated from equation 1 and Ct is the critical threshold for fluorescence.

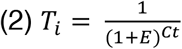

Then for each individual we summed the opsin gene expression across the four opsin genes and calculated the proportion of total expression that each gene exhibited.

Amplicons from the RT-qPCR for each gene (primer pair) were sequenced from one individual. Sanger sequencing of the amplicons was done at the NAPS Sequencing Centre at the University of British Columbia

